# Continuous tracking of task parameters tunes reaching control online

**DOI:** 10.1101/2022.06.13.495908

**Authors:** Antoine De Comite, Frédéric Crevecoeur, Philippe Lefèvre

## Abstract

A hallmark of human reaching movements is that they are appropriately tuned to the task goal and to the environmental context. This was demonstrated by the way humans flexibly respond to mechanical and visual perturbations that happen during movement. Furthermore, it was previously showed that the properties of goal-directed control can change within a movement, following abrupt changes in the goal structure. Such online adjustment was characterized by a modulation of feedback gains following switches in target shape. However, it remains unknown whether the underlying mechanism merely switches between prespecified policies, or whether it results from continuous and potentially dynamic adjustments. Here, we address this question by investigating participants’ feedback control strategies in presence of various changes in target width during reaching. More specifically, we studied whether the feedback responses to mechanical perturbations were sensitive to the rate of change in target width, which would be inconsistent with the hypothesis of a single, discrete switch. Based on movement kinematics and surface EMG data, we observed a modulation of feedback response clearly dependent on dynamical changes in target width. Together, our results demonstrate a continuous and online transformation of task-related parameters into suitable control policies.

**Significance statement:** Humans can adjust their control policy online in response to changes in the goal structure. However, it was unknown whether this adjustment resulted from a switch between two policies, or from dynamic and continuous adjustments. To address this question, we investigated whether online adjustments were tuned to dynamic changes in goal target which varied at different rates. Our results demonstrated that online adjustments were tuned to the rate of change in target width, suggesting that human reaching control policies are derived based on continuous monitoring of task-related parameters supporting online and dynamic adjustments.

## Introduction

Humans can execute reaching movements in various environments in the presence of unexpected disturbances such as visual or mechanical perturbations, that can interfere with their ability to succeed. Indeed, a large body of work characterized human control policies during reaching in presence of step mechanical (Cross et al. 2019; Knill et al. 2011; Lowrey et al. 2017; Nashed et al. 2012), visual (Georgopoulos et al. 1981; Prablanc and Martin 1992; Sarlegna and Mutha 2015; Soechting and Lacquaniti 1983), or vestibular perturbations (Keyser et al. 2017; Oostwoud Wijdenes et al. 2019). Crucially, the perturbations used in these experiments recruited feedback circuits without altering the limb dynamics, which allowed establishing the dependency of the control policy on task requirements. These results highlighted that reaching control policies flexibly adapted to a wide variety of contexts while relying on different sensory modalities.

To capture this feature, the control of upper limb reaching movements can be modeled in the framework of Optimal Feedback Control (OFC). This theory posits that reaching control policies optimize a performance index captured by a cost-function consisting of a weighted combination of motor cost and state-dependent movement penalties. This cost-function encompasses the task requirements by determining how to control the limb optimally with respect to this goal (Todorov 2004; Todorov and Jordan 2002). OFC has been used to model a diverse set of perturbation paradigms and established the flexibility of goal-directed feedback control in humans (Diedrichsen 2007; Diedrichsen and Dowling 2009; Izawa and Shadmehr 2008; Nashed et al. 2014; Omrani et al. 2013; Scott 2016).

It is important to realize that in most studies, the planning and control phases have been dissociated. Indeed, it was often assumed that the movement goal is selected prior to executing the corresponding control policy (Wong et al. 2015). In the OFC framework, the dissociation of planning and execution corresponds to the assumption that the feedback gains, and therefore the control policy, are derived prior to movement. In this view, it is unclear whether and based on which variables can the nervous system update control of an ongoing movement following changes in task-related parameters altering the movement goal, thereby implying a novel cost and requiring an adjustment of the policy.Crucially, we must distinguish perturbations as target jumps or mechanical loads, which computationally can be handled by altering the state vector without changing the controller, from changes in task requirements such as the structure of the target, that impose a change in the controller itself.

We recently demonstrated that the goal-directed policy used during reaching was adjusted online in response to changes in target width (De Comite et al. 2021). Here, we sought to investigate whether such adjustments reflected participants’ ability to switch between two prespecified control strategies, or whether they resulted from a feedback system considering continuous changes in the goal structure, and responded accordingly.

We addressed this question in two experiments where participants had to perform reaching movements towards a target the width of which could gradually decrease at different rates during movement, corresponding to a continuous modification of the target redundancy along its main axis. Two alternative hypotheses can be formulated: if adjustments in control policy do not integrate the dynamical changes in target, we expect to see stereotyped switches in behavior and feedback responses across conditions reflecting switches between two extreme cases (corresponding to maximal and minimal target widths). On the contrary, if dynamic changes are monitored, different rates of changes in target width should evoke different amounts of modulation in feedback responses. In agreement with the second alternative, we observed across the two experiments that participants adjusted their response to the rate of change in target width Together, our results demonstrate the existence of a feedback mechanism conveying continuous information about task-parameters and adjusting the control policies dynamically.

## Methods

### Participants

A total of 24 right-handed participants were recruited for this study and were enrolled in one of the two experiments. Fourteen participants (10 females) ranging in age from 18 to 30 years old took part to Experiment 1. The second group performed Experiment 2 and included 10 right-handed participants (5 females) ranging in age from 19 to 27 years old. Participants were naïve to the purpose of the study, had normal or corrected vision, and had no known neurological disorder. The ethics committee of the local university approved the experimental procedures and participants provided their written informed consent prior to the experiment.

### Experimental paradigm

Participants were seated on an adjustable chair in front of a Kinarm end-point robotic device (KINARM, Kingston, ON, Canada) and grasped the handle of the right robotic arm with their right hand. The robotic arm allowed movements in the horizontal plane and direct vision of both the hand and the robotic arm was blocked. Participants sat such that, at rest, their arm was approximately vertical and their elbow formed an angle of about 90°.Their forehead rested on a soft cushion attached to the frame of the robot. A semi-transparent mirror, located above the handle and reflecting a virtual reality display (VPixx, 120Hz) allowed participants to interact with visual targets. A white dot of 0.5cm radius aligned to the position of the right handle was displayed throughout the whole experiment.

### Experiment 1

In this experiment, participants (N=14) were instructed to perform reaching movements to a visual target initially represented as a wide rectangle (30×2.5cm) located 20 cm away from the home target in the y-direction. The home target was a circle of 1.5cm of diameter. The main axis of the rectangle was aligned with the x-axis and was orthogonal to the straight-line path from the home target to the center of the goal target (see A). Participants first had to bring the hand-aligned cursor in the home target displayed as a red circle that turned green as they reached it. After a random delay (uniform, between 1 and 2 s), the goal target was projected as a gray rectangle and participants could begin their movement whenever they wanted. There was no constraint on the reaction time. The exit from the home target was used as an event to determine reach onset, and starting then participants had to complete their movement between 350 and 600ms to successfully complete the trial. The trial was successfully completed if (i) they reached the goal target within the prescribed time window and (ii) they were able to stabilize the cursor in it for 500ms. The goal target turned green at the end of successful trials and red otherwise. To motivate the participants, a score corresponding to their number of successful trials was projected next to the goal target.

During movements, two types of perturbations could occur. The first one was a mechanical load consisting of a lateral step force applied by the robot to participants’ hand (53.6% of trials). The magnitude of this force was ± 9N aligned with the x-axis, with a 10-ms linear build-up. This force was triggered when the hand-aligned cursor crossed a virtual line parallel to the x-axis and located at 6cm from the center of the home target (see Figure1A, horizontal black dashed line). This step force was switched off at the end of the trial. The second type of perturbation was a visual change in target width starting when participants exited the home target (51.2% of trials). Hereafter we refer to the visual perturbation as the *target condition*. Participants had no information about the target condition prior to movement initiation. This change could either be an instantaneous change from a wide rectangle to a narrow square (*switch* condition, Figure1B, magenta) or a continuous change in target width either at a speed of −30cm/s (*slow* condition, green in Figure1B) or at a speed of −45.8cm/s (*fast* condition, blue in Figure1B). The speed of the *fast* condition was selected such that the target width at the end of the movement was similar to the *switch* conditions for the slowest correct movements. This was done to assess whether participants could anticipate the final width of the target and select a corresponding controller, which would produce identical responses in the *fast* and *switch* conditions. The decrease in target width stopped as participants entered the goal target. Importantly, the location of the center of the goal target did not change across conditions and was always aligned with the home target.

**Figure 1.**
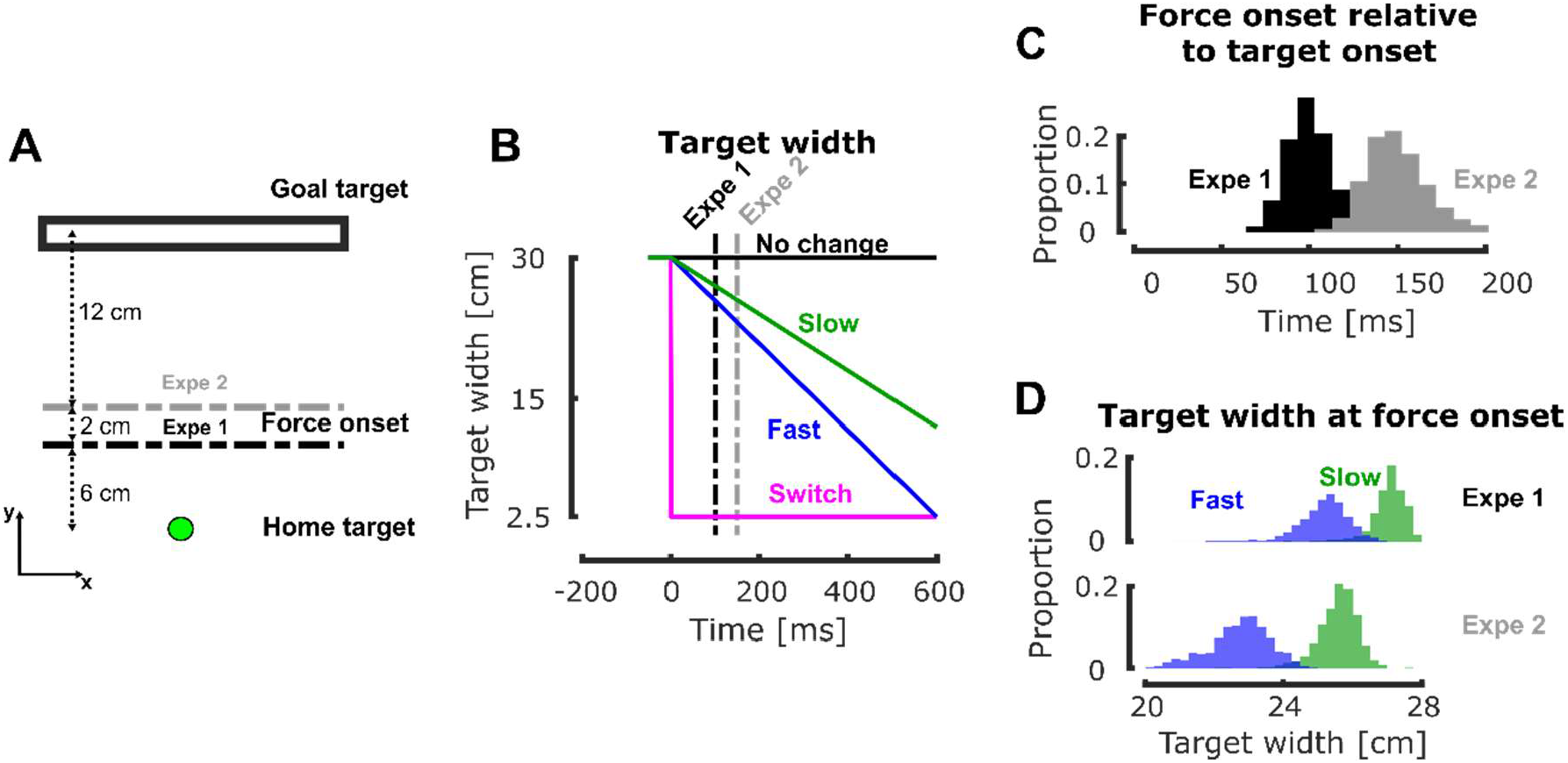
Experimental paradigms: **A** Schematic representation of the task paradigm. Participants had to perform reaching movement from the home target to the goal target, initially represented as a 30cm wide rectangle. During movement, they could experience mechanical step forces triggered in position at 6cm (Experiment 1, black line) or 8 cm (Experiment 2, gray line) from the home target and visual changes in target width (triggered as they exited the home target). **B** Evolution of the target width with respect to time in the different target conditions. The time axis is aligned on the visual perturbation onset defined by the onset of movement. The vertical dashed lines represent the median force onsets for Experiment 1 and 2. **C** Histograms of the distribution of the time interval between the visual and mechanical perturbation onset (resp. target and force onset) across all participants and conditions in Experiment 1 (black) and 2 (gray). **D** Histograms of the distribution of the target width at force onset for the two dynamical conditions (fast in blue and slow in green) across all participants in Experiment 1 (top) and 2 (bottom).

Unperturbed and perturbed trials were randomly interleaved such that participants could not predict the occurrence and the nature of disturbances. Participants were instructed to reach the target as it was actually displayed. They started with a 25-trials training block in order to become familiar with the task, the timing constraints, and the force intensity of perturbation loads. Crucially, this training block did not contain any visual perturbation. After completing this training block, participants performed 6 blocks of 82 trials. Each 82-trials block contained: 38 trials without mechanical perturbation (20 with no target change and 6 for each target condition) and 44 trials with mechanical perturbation (20 with no target change and 8 for each target condition, equally likely for rightward and leftward mechanical perturbations). Participants performed a total of 492 trials, including 24 of each combination of perturbed condition (direction of the mechanical perturbation and target condition). Participants were compensated for their participation.

**Table 1.** Trials distribution for each block of the two experiments.

|  | No change | Slow | Fast | Switch |
| --- | --- | --- | --- | --- |
| No | 20 | 6 | 6 | 6 |
| Left | 10 | 4 | 4 | 4 |
| Right | 10 | 4 | 4 | 4 |

### Experiment 2

We designed a second experiment which was a variant of the first one to assess reproducibility of the results in a slightly different version of the protocol, and also to investigate possible influence of the delay between the visual and mechanical perturbations on the modulation of feedback responses. Experiment 2 was almost identical to Experiment 1, except that the mechanical perturbation was triggered when the hand-aligned cursor crossed a virtual line parallel to the x-axis and located at 8cm (instead of 6) from the center of the home target (see Figure1A, grey dashed line). The intensity of this mechanical load was reduced compared to the main experiment (7N vs 9N) in order to keep a similar success rate. All the other experimental parameters (target conditions, number of trials and time constraints) were identical to those of Experiment 1.

Since both visual and mechanical perturbations were triggered based on position threshold (respectively when participants exited the home target and when they crossed a virtual line located at 6cm or 8cm from the center of the home target), there was some variability in the time span between these two perturbation triggers. The variability in this time span is represented in Figure1C (black and gray for Experiments 1 and 2 respectively) and had a median value of 96±6.41ms for Experiment 1 and 145±21.73ms for Experiment 2. As the target width in the *slow* and *fast* conditions were continuously changing with time, some variability was also present in the target width at the mechanical perturbation onset. In the *fast* condition we observed a median value of 25.6±0.3cm and 23.31±0.14cm while in the *slow* condition we observed a median value of 27.1±0.15cm and 26.3±0.54cm, respectively for Experiment 1 and 2, represented in Figure1D in blue (*fast*) and green (*slow*).

### Data collection and analysis

Raw kinematics data were sampled at 1kHz and low-pass filtered using a 4^th^ order double-pass Butterworth filter with cut-off frequency of 20 Hz. Hand velocity, acceleration and jerk were computed from numerical differentiation of the position using a 4^th^ order centered finite difference.

Surface EMG electrodes (Bagnoli surface EMG sensor, Delsys INC. Natick, MA, USA) were used to record muscles activity during movements. We measured the Pectoralis Major (Pect. Maj.) and the Posterior Deltoid (Post. Delt.) based on previous studies (Crevecoeur et al. 2019, 2020b; De Comite et al. 2021) showing in the same configuration that these muscles were stretched by the application of lateral forces, and therefore strongly recruited for feedback responses. Before applying the electrodes, the skin of participants was cleaned and abraded with cotton wool and alcohol. Conduction gel was applied on the electrodes to improve the quality of the signals. The EMG data were sampled at a frequency of 1kHz and amplified by a factor of 10 000. A reference electrode was attached to the right ankle of the participant. Raw EMG data from Pect. Maj. and Post. Delt. were band-pass filtered using a 4^th^ order double-pass Butterworth filter (cut-offs: 20 and 250Hz), rectified, aligned to force onset and averaged across trials or time as specified in the Results section. EMG data were normalized for each participant to the average activity collected when they maintained postural control against a constant force of 9N (rightward for Pect. Maj., leftward for Post. Delt.) This calibration procedure was applied after the second and the fourth blocks.

### Statistical analyses

Data processing and parameter extractions were performed using Matlab 2019a. We fitted linear mixed models (Brown and Prescott 2006) to infer the effect of target conditions on different kinematics parameters and on the EMG activities. These models were fitted using the *fitlme* function and the formula used was the following:

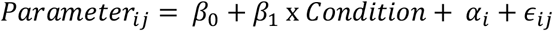

In this formula, the fixed predictors were the intercept (*β*) and the target condition (*β*_1_) while participants were included as a random offset (*α*_*i*_). The individual residual of trial *j* for participant *i*, captured by *ϵ*_*ij*_ followed a normal distribution. Each target condition was associated with an integer number such that they were ordered in decreasing order of constraints on the final target (*no change<slow<fast<*switch) and that positive/negative values for the regressor *β*_1_ indicate a decrease/increase of the measured parameter with the task difficulty. For these linear mixed model analyses that we performed, we reported the mean estimate of *β*_1_, its standard deviation, the t-statistics for this estimate and the corresponding p-value.

The continuous predictor for the condition can be seen as a non-linear transform of task difficulty and the parameter *β*_1_ can be interpreted as a slope, meaning that the more difficult the task is (with narrower target), the larger the feedback response. However, this approach can be criticised as the condition may also be considered as a categorical predictor. To address this concern, we also ran a discrete version of the linear mixed models where the target condition was defined as categorical. This categorical model confirmed the conclusion of the continuous one in all the conditions (results not shown). Post-hoc tests between pairs of target conditions were performed using similar linear mixed model applied on the two compared target conditions. For these post-hoc tests, we reported the mean estimate of *β*_1_, its standard deviation, the t-statistics for this estimate, the corresponding p-value, and the effect size defined as the standardized mean difference between two groups of independent observations (Lakens 2013).

In order to determine whether the timing of the mechanical perturbation relative to the onset of visual change could modulate the feedback responses, we compared the results of Experiments 1 and 2 as follows. We normalized the EMG activity by the intensity of the mechanical perturbations (9N and 7N for Exp. 1 and 2, respectively), and binned them within trial in the long-latency (LL, 50-100ms following perturbation onset) and early-voluntary (VOL, 100-180ms following perturbation onset) (Pruszynski et al. 2008; Pruszynski and Scott 2012). We then ran the following linear mixed effect models for each of these binned response value:

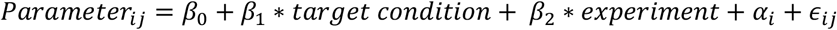

The fixed predictors are the intercept (*β*_*0*_), the target condition (*β*_1_) and the experiment (*β*_2_, a proxy for the onset of mechanical perturbation) while participants were included as random offset (*α*_*i*_). The individual residual of trial *j* for participant *i*, captured by *ϵ*_*ij*_ followed a normal distribution. As above, we verified that continuous and categorical definitions of the target condition yielded similar results and reported the statistics corresponding to the continuous predictor. Post-hoc tests between pairs of conditions were performed using linear mixed models applied on the two compared target conditions.

We also investigated a potential learning or habituation effect across *fast* and *slow* conditions as participants did not encounter those trials in the training phase. In order to investigate the lag between the first and last trials in the dynamical conditions (namely the *slow* and *fast* conditions that were not met during the training phase), we used a cross-correlation analysis applied on resampled data. We generated 1000 bootstrap samples from the individual acceleration profiles. For each of these samples, we computed the mean acceleration traces for the first and last trials and computed the cross-correlation between these two mean traces. We then extracted the peak value of this cross-correlation, corresponding to the lag between the two signals. The bootstrap resampling allowed us to obtain a distribution for this lag such that we could perform statistical analyses on it. Wilcoxon signed rank test was used to assess whether differences in lag were statistically different or not from zero.

In all our analyses, significance was considered at the level of p=0.05 even though we decided to exactly report any p-value that was larger than p=0.005 as previously proposed (Benjamin and Berger 2018).

## Results

### Experiment 1

Participants were asked to perform reaching movements to a target that was initially a 30cm wide rectangle, in all cases. During movement and in a random subset of trials, the target could either instantaneously turn into a 2.5cm wide square target (*switch* condition) or gradually decrease in width either at a high (*fast* condition) or low speed (*slow* condition). Additionally, unexpected mechanical perturbations were used during movements to evoke rapid motor responses and investigate their properties in relation with the change in target width

#### Kinematics

We observed that the target condition clearly influenced participants’ behavior. Indeed, the mean hand path trajectories in the mechanically perturbed conditions (Figure2A) differed across conditions. Consistent with our previous findings (De Comite et al. 2021) we observed online adjustments in the behavior in the *switch* condition (magenta) compared to the *no change* condition (black). These adjustments consisted of smaller lateral deviations in the *switch* condition (Figure2B and C, black and magenta traces). Interestingly, the behaviors in the dynamical conditions (*slow* and *fast*, green and blue respectively) differed from both the *no change* and *switch* conditions. In order to quantify these differences, we investigated the maximal hand deviation induced by the mechanical perturbations and the final hand position defined as the x-position of the hand as its velocity dropped below 2cm/s.

**Figure 2.**
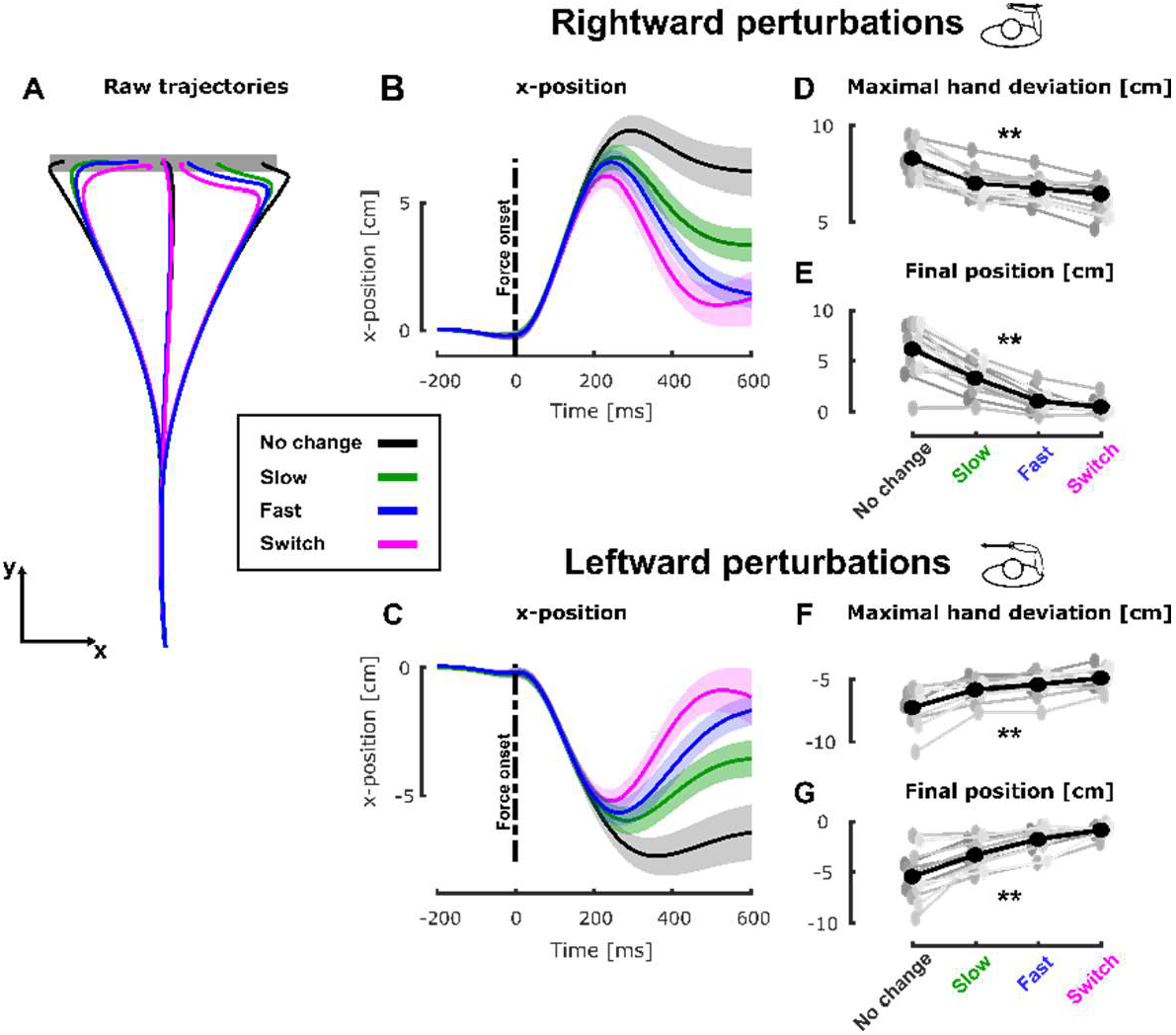
Experiment 1, Hand Kinematics during movement: **A** Group mean of the hand path for unperturbed and perturbed trials in the No Change (black), Slow (green), Fast (blue) and Switch (magenta) conditions. **B** Group mean and SEM of the x-position of participants’ hand as a function of time (aligned on force onset) for trials perturbed with rightward mechanical perturbations in the four target conditions. The black dashed line represents the onset of the mechanical perturbation. **C** Group mean and SEM of the x-position of participants’ hand as a function of time for trials perturbed with leftward mechanical perturbations in the four target conditions. The black dashed line represents the onset of the mechanical perturbation. **D** Group mean (black) and individual means (gray) of the maximal hand deviation in presence of a rightward perturbation for the four target conditions. **E** Group mean (black) and individual means (gray) of the final position for trials with rightward perturbation for the four target conditions. **F** Group mean (black) and individual means (gray) of the maximal hand deviation in presence of a rightward perturbation for the four target conditions. **G** Group mean (black) and individual means (gray) of the final position for trials with rightward perturbation for the four target conditions. ** p<0.005

The maximal lateral hand deviation induced by rightward mechanical perturbations (Figure2D) varied significantly across the target conditions. A linear mixed model (see Methods) revealed a significant effect of target condition (*β*_1_=0.0417±0.0022, t=18.14, p<0.005) on the maximal hand deviation with larger deviations for slower changes in target width. Post-hoc pairwise analyses revealed that the maximal hand deviation was larger in the *no change* condition than in *slow* and *fast* conditions (*slow* β_1_=-0.006±0.0006, t=-10.46, p<0.005, d= 0.60 and *fast* β_1_=-0.007±0.0006, t=-12.50, p<0.005, d=0.71). The hand deviation was larger in these dynamical conditions than it was in the *switch* condition (*fast* β_1_=-0.0021±0.0008, t=-2.56, p<0.005, d=0.11 and *slow* β_1_=-0.0017±0.0008, t=-3.15, p<0.005, d=0.21). Finally, we even observed that the hand deviation was larger in the *slow* than in the *fast* condition (β_1_=-0.0013±0.0006, t=-2.14, p=0.0319, d=0.1527). Similar results were observed for leftward mechanical perturbations (see Figure2F, linear mixed models: *β*_1_=0.0033±0.0002, t=16.2, p<0.005).

Similarly, we observed that the final hand position along the x-axis, computed as the hand position when the total velocity dropped below 2cm/s, exhibited similar dependency on the target condition. Indeed, a linear mixed model analysis (see Methods) revealed a significant effect of the target condition (*β*_1_=-0.008±0.0003, t=-25.75, p<0.005). As for the maximal hand deviation, post-hoc pairwise analyses revealed that both dynamical conditions were characterized by less eccentric final hand positions than the *no change* condition (*slow*, β_1_=-0.014±0.0008, t=-16.37, p<0.005, d=0.80 and *fast* β_1_=-0.025±0.0008, t=-28.74, p<0.005, d=1.27). These final hand positions in the *slow* condition were more eccentric than the one in the *switch* condition, no differences were found between the *fast* and *switch* conditions (*fast* β_1_=-0.0018±0.0013, t=1.36, p=0.17, d=0.09 and *slow* β_1_=-0.008±0.0013, t=-6.49, p<0.005, d=0.43). The final hand positions in the *slow* condition were significantly more eccentric than those in the *fast* condition (β_1_=-0.01±0.0008, t=-13.10, p<0.005, d=0.83). Trials that included a leftward mechanical perturbation (see Figure2G) contained the same effects (linear mixed models : *β*_1_=0.008±0.0003, t=27.89, p<0.005).

#### Muscle activity

The kinematics results that we reported indicated that participants were able to adjust their control strategy during movements according to dynamical changes in movement goal. They were even able to tune their adjustment to the speed of these dynamical changes. We hypothesized that the stretched EMG activity in Pectoralis Major (Pect. Maj.) and Posterior Deltoid (Post. Delt.) should also depend on the target condition. If such modulation exists in the long-latency epoch, 50-100ms following the onset of the mechanical perturbation, it would indicate that the adjustment in behavior did not only reflect changes in voluntary intent but also changes in reflexive responses previously associated with goal-directed state-feedback control (Crevecoeur and Kurtzer 2018; Pruszynski and Scott 2012).

We observed that the target condition modulated the EMG activity of the muscles stretched by the mechanical perturbation. Figure3A and B represent the mean EMG activities collapsed across participants for trials perturbed by rightward or leftward perturbation in all target conditions in the stretched (full lines) and shortened muscles (dashed lines). Visual inspection of target specific responses for the stretched muscles, obtained by subtracting the *no change* condition, confirmed this modulation of the EMG response (see Figure3C and D respectively for Pect. Maj. and Post. Delt.). In order to characterize this modulation, the EMG activity of the stretched muscle was averaged in the long latency (LL 50-100ms after force onset) and early-voluntary time epochs (VOL 100-180 ms after force onset) for each perturbation direction. The deviations from the mean activity in these time bins are reported in Figure3E and F for stretched Pect. Maj. in the LL and VOL windows at population (black) and individual (gray) levels.

**Figure 3.**
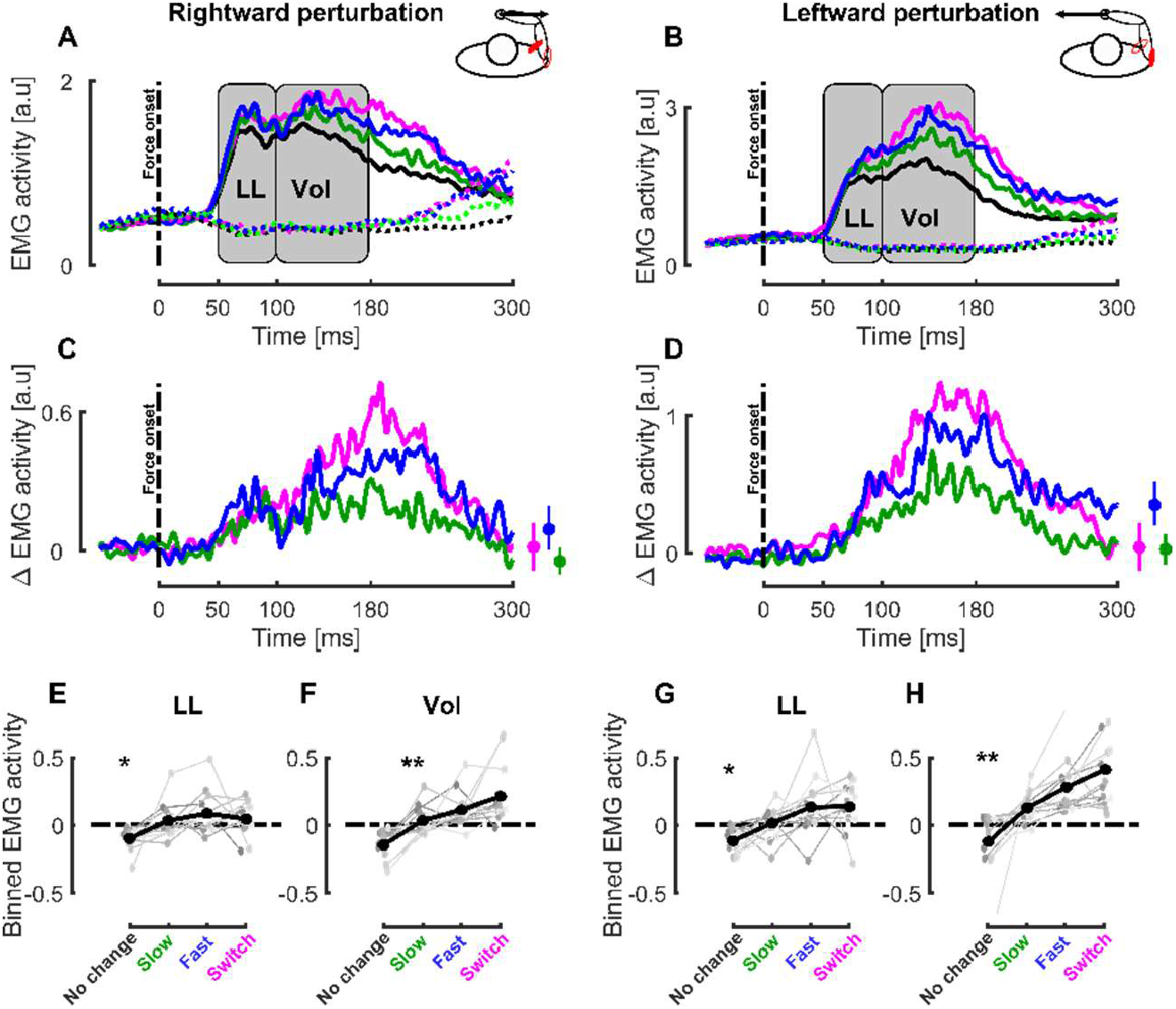
Experiment 1, EMG activity : **A** Group mean for the stretched (Pectoralis Major, full lines) and shortened (Posterior Deltoid, dashed lines) responses to rightward perturbations in the different target conditions. The gray rectangles represent the long latency (LL) and voluntary (VOL) epochs where the EMG activity was averaged to perform statistical analyses. The black dashed line represents mechanical perturbation onset and the time axis is aligned with force onset. **B** Group mean for the stretched (Posterior Deltoid, full lines) and shortened (Pectoralis Major, dashed lines) responses to leftward perturbations in the different target conditions. **C** Group mean of target-specific EMG responses to perturbation for Pectoralis Major in the presence of rightward perturbation for the switch (magenta), fast (blue) and slow (green) target conditions. Time axis is aligned with force onset. The small insets represent the mean and SEM target-specific EMG responses at the end of the movement. **D** Group mean of target-specific EMG responses to perturbation for Posterior Deltoid in the presence of leftward perturbation for the switch (magenta), fast (blue) and slow (green) target conditions. Time axis is aligned with force onset. The small insets represent the mean and SEM target-specific EMG responses at the end of the movement. **E-F** Group mean (black) and individual means (gray) of the binned EMG activity in the LL (E) and VOL (F) time windows for Pectoralis Major in the presence of rightward perturbations for the different target conditions. **G-H** Group mean (black) and individual means (gray) of the binned EMG activity in the LL (G) and VOL (H) time windows for Posterior Deltoid in the presence of leftward perturbations for the different target conditions. * p<0.05, ** p<0.005

Strikingly, we observed a significant effect of target condition on the modulation of the Pect. Maj. response in the LL (linear mixed models : *β*_1_=-0.029±0.005, t=-5.70, p<0.005) and VOL window (linear mixed models : *β*_1_=-0.060±0.00495, t=-12024, p<0.005), respectively represented in Figure3E and F. These negative values indicated larger responses for faster changes in target width. To further investigate these differences, we performed pairwise post-hoc comparisons between the different target conditions using linear mixed models (see Methods). In the LL window, we did not observe any difference between the different dynamical conditions (*switch/fast* : *β*_1_=0.02±0.021, t=0.96, p=0.33, d=0.05, *switch/slow β*_1_=-0.0038±0.013, t=-0.29, p=0.77, d=0.015 and *slow/fast β*_1_=-0.0525±0.0419, t=-1.25, p=0.21, d=0.064), even though they all differed from the *no change* condition (p<0.005 for all conditions). However, these pairwise comparisons revealed significant differences in the VOL time window between the dynamical conditions (*switch/fast β*_1_=-0.05±0.019, t=-2.50, p=0.012, d=0.11, *switch/slow β*_1_=-0.059±0.013, t=-4.35, p<0.005, d=0.21 and *slow/fast β*_1_=-0.079±0.04, t=-1.96, p=0.048, d=0.1).

The same modulation of the EMG activity with the target condition was observed in Post. Delt. for both LL (mixed models: *β*_1_=-0.046±0.007, t=-6.42, p<0.005 see Figure3G) and VOL time epochs (mixed models: *β*_1_=-0.015±0.008, t=-18.66, p<0.005 see Figure3F) when stretched by leftward perturbation. Interestingly, the pairwise post-hoc comparisons revealed significant differences between the dynamical conditions in both the LL (*switch/fast* : *β*_1_=-0.005±0.019, t=-0.03, p=0.97, d=0.002, *switch/slow β*_1_=-0.03±0.012, t=-2.336, p=0.019, d=0.14, and *slow/fast β*_1_=-0.1193±0.057, t=-2.09, p=0.036, d=0.12) and the VOL time window (*switch/fast* : *β*_1_=-0.072±0.021, t=-3.35, p<0.005, d=0.14, *switch/slow β*_1_=-0.12±0.016, t=-7.33, p<0.005, d=0.33 and *slow/fast β*_1_=-0.25±0.058, t=-4.33, p<0.005, d=0.19). These differences indicated that both reflexive and voluntary responses were modulated by the dynamical change in target width, and suggest that they were even tuned to the rate of change in target width. The significance of the post-hoc effect between the *switch/fast* conditions in the voluntary epochs rules out the possibility that participants only used the predicted final target width to modulate their behavior.

Altogether, these results indicate that participants adjusted their behavior during movements in response to dynamical changes in target shape. Indeed, we showed that the hand deviation induced by the mechanical perturbations was different in the dynamical (*slow* and *fast*) and in the static conditions (*no change* and *switch*). Moreover, we reported larger hand deviation for the *slow* than for the *fast* condition: indicating that the rate of change in target width was integrated in the control strategy. The differences observed in acceleration profiles and EMG correlates confirmed this finding. The sensitivity of the online adjustments of control policy to dynamical changes and speed of changes suggest the existence of a mechanism able to finely tune to control strategies within movement.

### Experiment 2

Although there was, in Experiment 1, a significant effect in the long-latency window, the pairwise comparisons did not allow to conclude that the modulation was as gradual as in the VOL epoch. We designed Experiment 2 to test the possibility that the shallower modulation in the LL, compared to that in the VOL epoch (see panels E vs F and G vs H), was due to a too short delay between the onset of visual changes and the mechanical perturbation, which could therefore leave too little time to develop a clear modulation in the long-latency epoch. In Experiment 2, the onset of the visual perturbation was similar as in Experiment 1 but that of the mechanical perturbations occurred later (150ms after the visual onset instead of 100ms in Experiment 1) which allowed more time to adjust control policies as a delay of 150ms was previously reported between the onset of change in target width and changes in control (De Comite et al. 2021).

The impact of the different target conditions on the behavior was qualitatively similar to that of Experiment 1 described in Figure2. We then investigated the modulation of the EMG activity during Experiment 2 in the LL and VOL epochs with the same linear mixed model as in Experiment 1. We observed significant modulation in both the LL (Pect. Maj.: *β*_1_=-0.021±0.04, t=-4.43, p<0.005 and Post. Delt.: *β*_1_=-0.014±0.006, t=-2.09, p=0.036) and VOL (Pect. Maj.: *β*_1_=-0.041 ±0.006, t=-6.17, p<0.005 and Post. Delt.: *β*_1_=-0.062±0.011, t=-5.83, p<0.005) time epochs during this control experiment. Similar to what was found in Experiment 1, we observed a shallower modulation in the LL time epoch than in the VOL epoch indicating that the design of Experiment 1 did not unintentionally reduce the modulation of the response in the LL time epoch.

We investigated whether the differences in responses observed between the *fast* and the *slow* conditions result from differences in rates of change in target width or from the instantaneous target width at perturbation onset, by comparing the normalized EMG responses observed in Experiments 1 and 2. If the hypothesis whereby these modulations of the feedback responses are mediated by the width of the target at perturbation onset holds, we should observe larger responses in Experiment 2 as the mechanical perturbations were triggered later resulting in smaller target width at perturbation onset (see Figure4A and B). To test this hypothesis, we grouped the normalized stretch muscle activities, binned in the LL and VOL time epochs, from both experiment and investigated a potential effect of the experiment (see Methods). We did not observe any differences between the normalized responses of Experiment 1 and 2, neither in the LL epoch (*β*_2_ = 0.12 ± 0.08, t=1.58, p=0.1134, Figure 4-D) nor in the VOL epoch (*β*_2_ = 0.17 ± 0.11, t=1.43, p=0.1516, Figure 4-F). Since we did not observe differences between the feedback responses across the two experiments, we decided to pool these responses to gain a more robust statistical description of the main effect in the LL and VOL time windows. We grouped the muscle activity of the stretched muscles from both experiments (see Figure 4G for the mean traces) and used linear mixed models that considered target conditions and experiments as fixed factors (see Methods). We found a main effect of the target condition in both LL (*β*_1_=-0.03±0.003, t=-9.62, p<0.005) and VOL (*β*_1_=-0.09±0.003, t=-25.61, p<0.005) epochs. Post-hoc pairwise comparisons performed between conditions in these two time epochs also reported differences demonstrating larger feedback responses for more constrained movements (LL: *switch vs fast β*_1_*=-0*.*024±0*.*029, t=-0*.*89, p=0*.*39*, d=0.02 *switch vs slow β*_1_*=-0*.*039±0*.*013, t=-2*.*80, p<0*.*005, d=0*.*10* and *fast vs slow β*_1_*=-0*.*063±0*.*028, t=-2*.*24, p=0*.*005*, d=0.07 VOL: *switch vs fast β*_1_*=-0*.*24±0*.*035, t=-7*.*04, p<0*.*005, d=0*.*35,switch vs slow β*_1_*=-0*.*19±0*.*017, t=-11*.*504, p<0*.*005, d=0*.*21* and *fast vs slow β*_1_*=-0*.*15±0*.*031, t=-4*.*89, p<0*.*005, d=0*.*14*).

**Figure 4.**
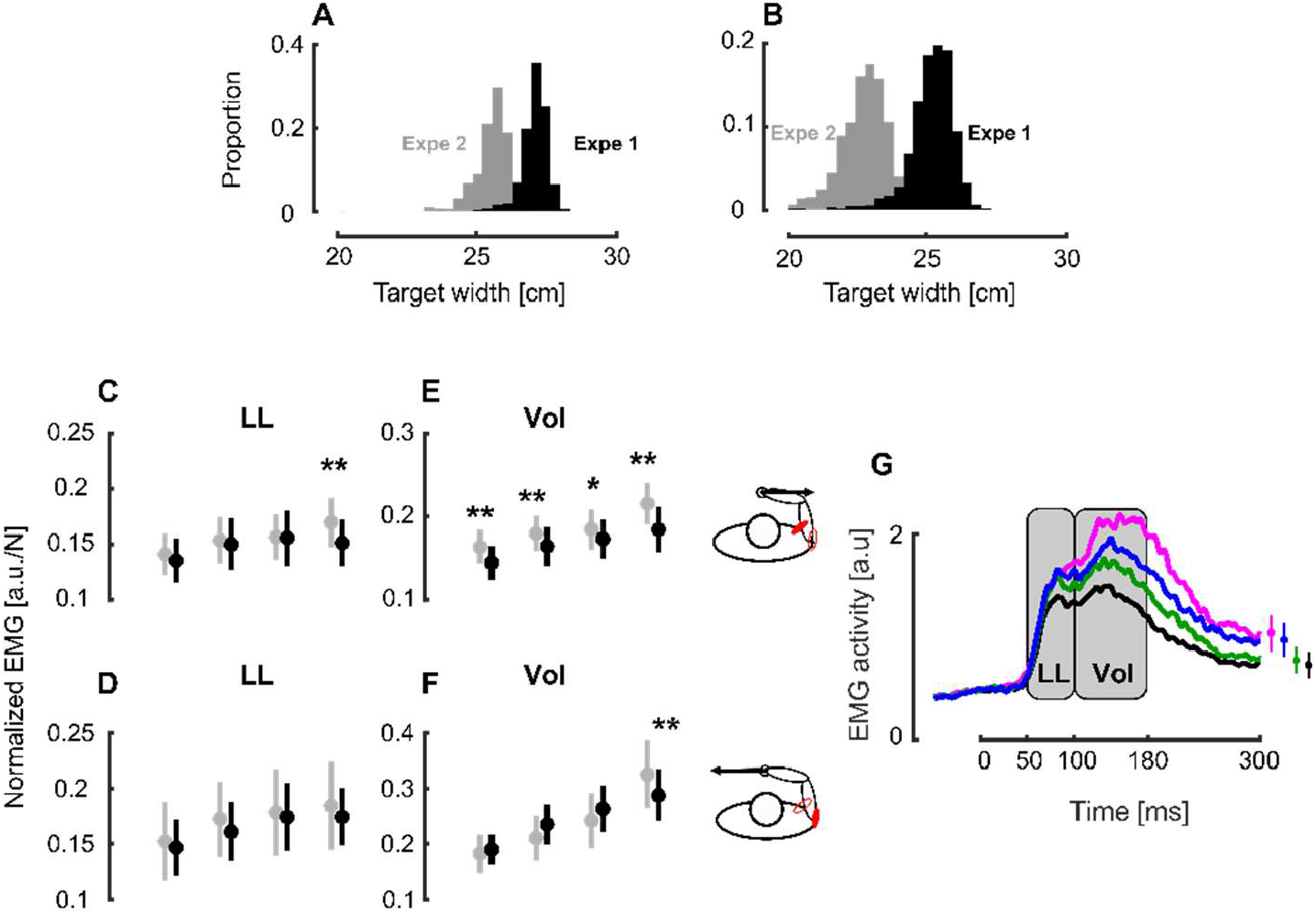
Normalized EMG across experiments: **A** Distribution histogram of the target width at perturbation onset in the slow target condition in Experiment 1 (black) and 2 (grey). **B** Distribution histogram of the target width at force onset in the fast target condition in Experiment 1 (black) and 2 (grey). **C** Group mean and SEM collapsed across participants of the normalized Pect. Maj. EMG activities in the LL time window in Experiment 1 (black) and 2 (gray) across conditions in presence of rightward perturbations. **D** Group mean and SEM collapsed across participants of the normalized Post. Delt. EMG activities in the LL time window in Experiment 1 (black) and 2 (gray) across conditions in presence of leftward perturbations. **E** Group mean and SEM collapsed across participants of the normalized Pect. Maj. EMG activities in the VOL time window in Experiment 1 (black) and 2 (gray) across conditions in presence of rightward perturbations. **F** Group mean and SEM collapsed across participants of the normalized Post. Delt. EMG activities in the VOL time window in Experiment 1 (black) and 2 (gray) across conditions in presence of leftward perturbations. **G** Group mean across participants, experiments and muscles of the stretched muscle activity for the different target conditions. The gray rectangles represent the long-latency and voluntary time epochs. The time axis is aligned on force onset and the insets at the right of the panel represent the mean and SEM of the stretched muscles activity at the end of movement. * p<0.05, ** p<0.005

This second experiment revealed that the modulation of EMG activity in the long-latency epoch was small but robust and reproducible. We also found across the two experiments, for which the target width at perturbation onset was different, that the responses were very similar. Observe that the perturbation in Experiment 2 were triggered a bit later, which potentially increase the response gains (Poscente et al. 2021). Thus, this effect should add to a potential sensitivity to target width. Nevertheless, we found essentially similar normalized EMG in spite of (slightly) later occurrence and smaller instantaneous width. This result suggests that the underlying neural pathways may consider the speed or rate of change of target width, which is clearly consistent with our hypothesis that continuous change in task parameters modulate control gains dynamically.

### Differences between the first and last trials in dynamical conditions

Interestingly, we observed that participant’s behaviour during the *fast* and *slow* conditions changed across blocks. Figure5A and B represents the mean and SEM of the position along the x-axis for the first (full line) and last trials (dashed line) in the *fast* condition for rightward and leftward mechanical perturbations respectively. We observed that these first and last trials differed and decided to take a look at their acceleration profiles in order to quantify these differences. The corresponding acceleration profiles are represented in Figure5C and D. We observed a consistent and significant lag of the last trial with respect to the first one. This lag was computed by taking the median of the lags distribution that was obtained from the maximal values of the cross-correlation between the first and last *fast* trials of the fourteen subjects with 10 000 bootstrap samples (see Methods). The resulting distribution of this lag, obtained through this resampling method is represented in Figure5E. This method revealed a median lag of −18ms (depicted with a blue vertical line in Figure5E) that was significantly smaller than zero (signrank test z=-32.18, p<0.005).

**Figure 5.**
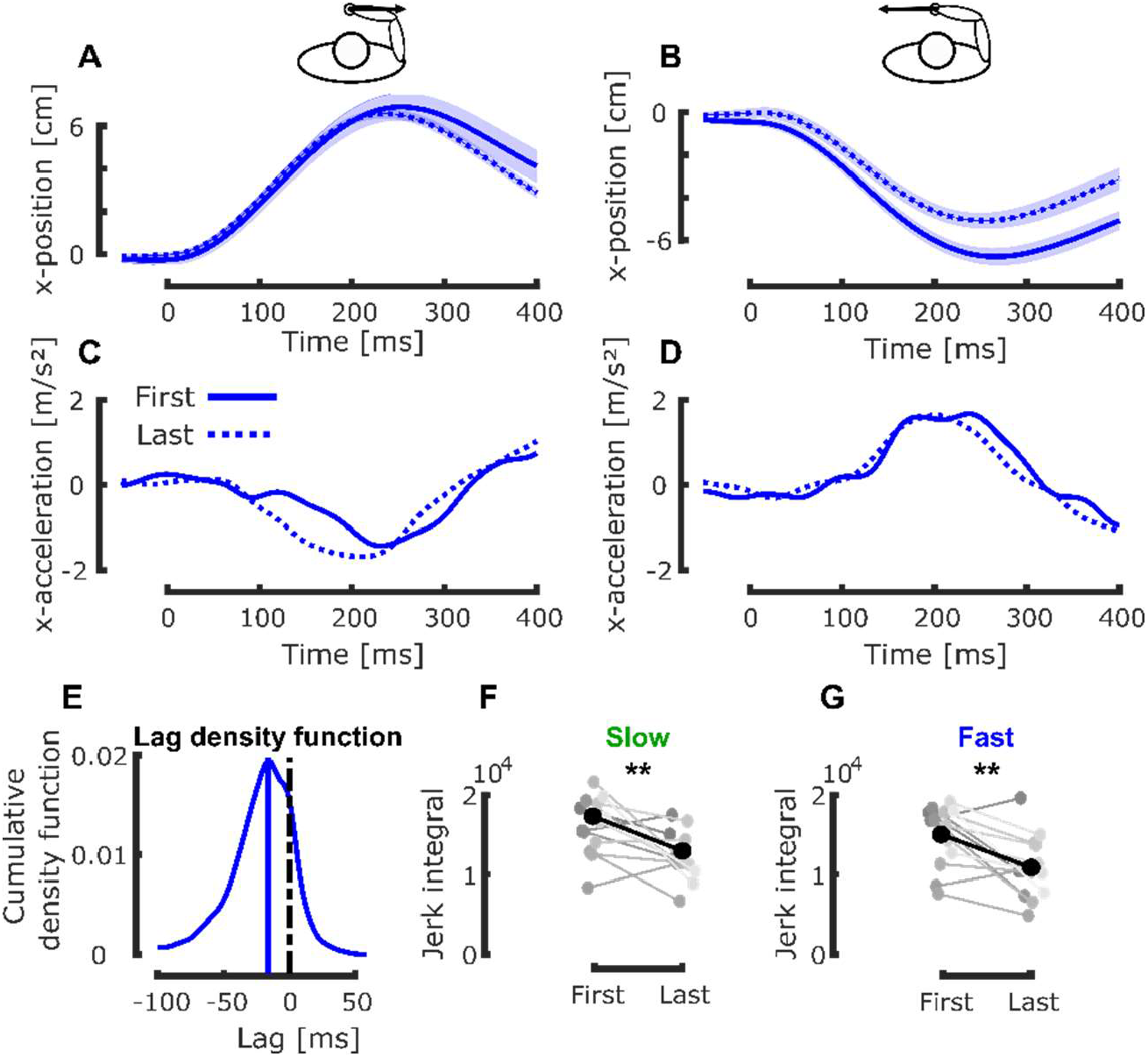
Between trials analyses : **A** Group mean and SEM collasped across participants of the first (full line) and last (dashed line) trials in the fast condition with rightward mechanical perturbation. Time axis is aligned on force onset. **B** Group mean and SEM collapsed across participants of the first (full line) and last (dashed line) trials in the fast condition with leftward mechanical perturbation. The black dashed line correponds to the difference between the first and last trials. Time axis is aligned on force onset. **C** Group mean of the acceleration profiles of the first (full line) and last (dashed line) trials in the fast condition with rightward mechanical perturbation. Time axis is aligned with force onset. **D** Group mean of the acceleration profiles of the first (full line) and last (dashed line) trias in the fast condition with leftward mechanical perturbation. Time axis is aligned with force onset. **E** Cumulative density function of the lag between the last and first acceleration profiles for both perturbation directions. The blue vertical line correpsonds to the median value. **F** Group mean (black) and individual means (gray) of the integral of the absolute value of the jerk for the first and last trials in the fast target condition. **G** Group mean (black) and individual means (gray) of the integral of the absolute value of the jerk for the first and last trials in the slow target condition. ** p<0.005

The first and last trials of each dynamical condition also differed in the smoothness of their acceleration profile as shown in Figure5C-D for the *fast* condition. This difference in smoothness was quantified by comparing the integral of the absolute values of the derivatives of these acceleration profiles: the jerk. We reported in Figure5F-G these integrals for all participants in the *slow* and *fast* conditions respectively. In the *fast* condition, the final state was less jerky than the first one as reported by a signrank test (z=-2.835, p<0.005). Similar results were obtained in the *slow* condition (signrank test z=-3.19, p<0.005) indicating an increase in the smoothness of the acceleration profiles.

Thus, there were measurable behavioral changes that could be related to practice, however they did not interfere with the interaction between target condition and behavior. Indeed, we still observed a significant modulation of the EMG activity in both LL (linear mixed models, *β*_1_=-0.041±0.009, t=-4.21, p<0.005) and VOL epochs (linear mixed models, *β*_1_=-0.144±0.014, t=-10.11, p<0.005) when we only considered the last twelve trials of each dynamical condition for all participants.

## Discussion

We investigated how humans responded to continuous changes in target width during reaching. More specifically, we studied participants’ behavior as they were reaching to a target, initially represented as a wide rectangle, with time varying width. We observed that the way participants responded to unexpected mechanical perturbations depended on the target condition and specifically on the rate of change in target width during movement. This demonstrated that the control policies used to perform reaching movements were adjusted online to the specific change in target width, which captures participants’ ability to continuously track and respond to task parameters during movement.

Here, we leveraged an experimental paradigm developed in a previous work (De Comite et al. 2021), consisting of abrupt changes in target structure within movements, to dynamically alter the task constraints and investigate whether participants’ control policies were adjusted online. This paradigm exploits the *minimum intervention principle* (Todorov and Jordan 2002) which states that participants only correct deviations that interfere with the task success during reaching movements. This means that participants exploit the target redundancy when available, even in the absence of perturbations (Berret et al. 2011; Knill et al. 2011; Nashed et al. 2012; Scholz et al. 2000; Togo et al. 2017; Vetter et al. 2002). The observed behaviour and the feedback responses to mechanical perturbations confirmed that these control policies were adjusted during movement. Indeed, we reported modulations induced by the different dynamical changes in target width, corresponding to different alterations of the cost-function and that this mechanism considered the rate of change in target width. In our view, these results demonstrate the existence of a mechanism that adjusts the control policy during movement thanks to a continuous tracking of target width. In the present study, this task-specific adjustment in control relied on visual processing of task-parameters and impacted long-latency and early voluntary responses to the mechanical perturbations.

We must emphasize a critical difference between a feedback response to an external perturbation and the results that we highlighted here. In standard perturbation paradigms, visual or mechanical events alter the state of the system, including limb and target position, velocity, and higher order derivatives. These perturbation paradigms allowed showing that the control policy used to perform movement is tuned to the task-goal as demonstrated by the goal-dependent characteristics of the feedback responses to disturbances (Keyser et al. 2017; Knill et al. 2011; Lowrey et al. 2017; Nashed et al. 2012; Sarlegna and Mutha 2015). These feedback responses are defined by rapid feedback loops (Figure6 inner loop, gray) whose latencies depend on the sensory modalities involved (Franklin and Wolpert 2008; Knill et al. 2011; Pruszynski and Scott 2012; Scott 2016). In the case of mechanical disturbances applied to the limb, this inner feedback loop is mediated by long-latency feedback pathways that have a latency of 50ms (Pruszynski and Scott 2012). Here, we probed not only the feedback responses to changes in the state of the system, but also the change in the controller itself in response to dynamic changes in task parameters during movement. Taken in the context of Optimal Feedback Control (OFC) (Scott 2004; Shadmehr and Krakauer 2008; Todorov and Jordan 2002), the task-parameters (such as target width) define the cost-function and the control law (Nashed et al. 2012), which is derived from these cost parameters. We demonstrated that this selection of the control policy based on the task parameters is itself continuous and must be considered in closed loop control models of human reaching movements (Figure6 outer loop, black). The latency of this outer loop was about 150ms as reported in previous work (De Comite et al. 2021), here we used this number to design the task, and highlighted that indeed dynamic changes in the task parameters have an impact in the long-latency feedback pathways.

**Figure 6.**
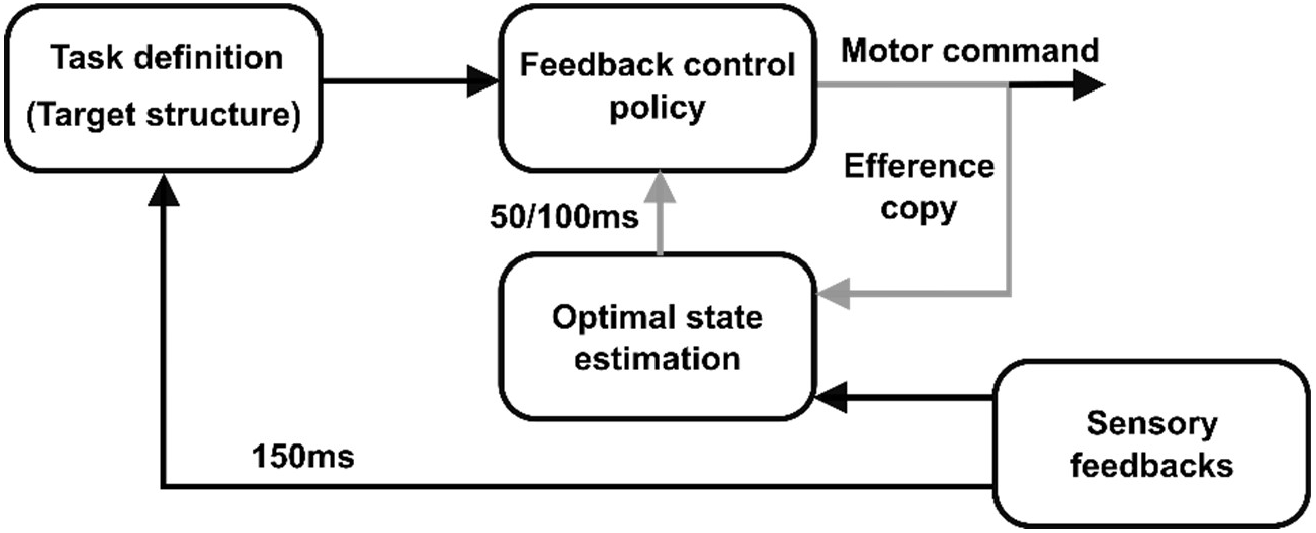
Modified goal-directed feedback control architecture. The inner loop (in gray), combining sensory feedbacks and efference copy is the one responsible for the goal-directed feedback responses observed behaviorally. The latency of this inner loop is 50-100ms depending on the sensory modalities involved. The outer feedback loop (in black) that modifies the task definition and the feedback control policy captures the online adjustments in control policy elicited by continuous alteration of the target structure during movement.

This interpretation implies a possible overlap of movement planning and execution, as participants may alter their motor plan during an ongoing movement. Such overlap of planning and execution has been suggested in studies reporting that reaction times prior to movement initiation could be shortened at the price of a reduced accuracy (Haith et al. 2016; Orban de Xivry et al. 2017). More recently, it has been suggested that such overlap of planning and execution could occur during movement in presence of visual perturbations (Cesonis and Franklin 2021, 2020; Dimitriou et al. 2013). However, these results could also be explained by other mechanisms such as an infinite horizon controller (Li et al. 2018). In our previous study, we provided clear evidence for this overlap of movement planning and execution in presence of perturbations that altered the cost-function from which the control policy is derived (De Comite et al. 2021). This overlap is also necessary to explain the present results as participants have to continuously adjust their control strategy in response to the change in target width.

These continuous adjustments of control policy are reminiscent of the theoretical framework of model predictive control (see (Lee 2011) for review). This framework posits that the control policy is continuously adjusted during movement in order to integrate any change in the cost-function or in the environmental context. An alternative hypothesis to explain these online adjustments in control policy is that participants switched between several pre-specified control policies. A similar process was suggested to account for the selection of the most appropriate strategy specified in parallel (Chapman et al. 2010; Gallivan et al. 2016, 2017; Wong and Haith 2017) but was questioned and compared to a single optimal intermediate motor plan (Alhussein and Smith 2021; Haith et al. 2015). Our experimental paradigm differed as multiple options were never presented at the same time and changes between targets occurred within movements.

We favour an interpretation that assumes dynamical adjustments of the control strategy even though we cannot formally rule out the possibility of discrete switches between different pre-defined controllers. However, there are observations that do plead for a continuous and dynamic monitoring. First, we observed that the first *slow* and *fast* trials where different, although participants had not encountered these conditions during training and therefore could not have acquired a controller tuned to these specific conditions at that time. This was observed in spite of the fact that these first trials were jerkier than the later ones, which could indicate that even when participants had not familiarized with the dynamical conditions, they seemed to exploit well the outer feedback loop as input to the controller relative to target structure (Figure 6, outer loop). One caveat to the hypothesis of continuous monitoring was that the normalized feedback responses across Experiments 1 and 2 (Figure 4) did not change much with longer viewing time, which suggests that there may be constraints on the amount of modulation that can take place. Nevertheless, this absence of modulation of the feedback responses within movement must be interpreted with caution because other factors such as time or urgency also modulate these feedback responses within movement (Crevecoeur et al. 2013; Dimitriou et al. 2013; Poscente et al. 2021).

A parallel can be drawn between the present study and a series of studies that reported within-trials tuning of feedback corrections when exposed to velocity-dependent force fields randomly (Crevecoeur et al. 2020a, 2020b; Mathew et al. 2020; Mathew and Crevecoeur 2021). It proposed that continuous tracking of model parameter also happens during movement, suggesting that adaptation to an altered plant dynamics also happens online (Crevecoeur et al. 2020b). In the present study, we reported the online tracking of cost parameters that define the movement goal. Interestingly, besides the conceptual similarity between these two processes linked to online evaluation of task or dynamical parameters, they were associated to different latencies: ∼150ms for updating the control policies following changes in movement goal (based on the present and on (De Comite et al. 2021)), while a latency of ∼250ms was associated with the online tuning of the feedback controller (Crevecoeur et al. 2020b). It is therefore conceivable that they engage dissociable neural operations that remain to be investigated.

An interesting question is to determine which neural substrates are involved in the online modulation of the reaching controller. Sensorimotor control is mostly supported by multiple cortical areas, the basal ganglia, and the cerebellum (Haar and Donchin 2020; Scott 2016; Shadmehr and Krakauer 2008). We identify two neural pathways that likely underlie the online changes in behavior documented in the present study. The first is that the parametric feedback controller supported by the long-latency feedback (Crevecoeur and Kurtzer 2018; Pruszynski and Scott 2012) is modulated online. Such modulation must have occurred based on visual input which has a fast route to the network supporting long-latency responses through associative areas in the parietal cortex (Cross et al. 2021). The second pathway that is likely involved is related to the definition of the task demands, which includes the basal ganglia known to represent motor costs (Mazzoni et al. 2007; Shadmehr and Krakauer 2008). Since our task paradigm altered the motor costs by modulating the target width, it is conceivable that the adjustment in control policy depended on a signal originating from the basal ganglia inducing the change of controller or a selection of a different controller. Note that this cannot happen independently of the visual input, and thus any interaction between rapid feedback pathways conveying information about the target structure, and selection of controllers based on a representation of motor costs in BG may support the observed change in behavior.

To sum up, we reported here that humans are able to dynamically adjust their control policy when they experience a dynamical change in task demands. These findings highlight the existence of a continuous monitoring of task-related parameters which supports dynamic changes in online feedback control.

## Notes

### Competing Interest Statement

The authors have declared no competing interest.

## References

Alhussein L, Smith MA. Motor planning under uncertainty. eLife 10: e67019, 2021.

Berret B, Chiovetto E, Nori F, Pozzo T. Manifold reaching paradigm: How do we handle target redundancy? J Neurophysiol 106: 2086–2102, 2011.

Brown H, Prescott R. Applied mixed models in medicine. 2nd ed. Chichester, England; Hoboken, NJ: John Wiley, 2006.

Cesonis J, Franklin D. Mixed-horizon optimal feedback control as a model of human movement. arXiv, 2021.

Cesonis J, Franklin DW. Time-to-target simplifies optimal control of visuomotor feedback responses. eNeuro 7: 514–519, 2020.

Chapman CS, Gallivan JP, Wood DK, Milne JL, Culham JC, Goodale MA. Reaching for the unknown: Multiple target encoding and real-time decision-making in a rapid reach task. Cognition 116: 168–176, 2010.

Crevecoeur F, Kurtzer I. Long-latency reflexes for inter-effector coordination reflect a continuous state feedback controller. J Neurophysiol 120: 2466–2483, 2018.

Crevecoeur F, Kurtzer I, Bourke T, Scott SH. Feedback responses rapidly scale with the urgency to correct for external perturbations. J Neurophysiol 110: 1323–1332, 2013.

Crevecoeur F, Mathew J, Bastin M, Lefevre P. Feedback adaptation to unpredictable force fields in 250ms. eNeuro 7, 2020a.

Crevecoeur F, Scott SH, Cluff T. Robust Control in Human Reaching Movements: A Model-Free Strategy to Compensate for Unpredictable Disturbances. J Neurosci 39: 8135–8148, 2019.

Crevecoeur F, Thonnard J-L, Lefevre P. A very fast time scale of human motor adaptation: within movements adjustments of internal representations during reaching. eNeuro 7, 2020b.

Cross KP, Cluff T, Takei T, Scott SH. Visual Feedback Processing of the Limb Involves Two Distinct Phases. J Neurosci 39: 6751–6765, 2019.

Cross KP, Cook DJ, Scott SH. Convergence of proprioceptive and visual feedback on neurons in primary motor cortex. Neuroscience.

De Comite A, Crevecoeur F, Lefèvre P. Online modification of goal-directed control in human reaching movements. J Neurophysiol 125: 1883–1898, 2021.

Diedrichsen J. Optimal Task-Dependent Changes of Bimanual Feedback Control and Adaptation. Curr Biol 17: 1675–1679, 2007.

Diedrichsen J, Dowling N. Bimanual coordination as task-dependent linear control policies. Hum Mov Sci 28: 334–347, 2009.

Dimitriou M, Wolpert DM, Franklin DW. The Temporal Evolution of Feedback Gains Rapidly Update to Task Demands. J Neurosci 33: 10898–10909, 2013.

Franklin DW, Wolpert DM. Specificity of Reflex Adaptation for Task-Relevant Variability. J Neurosci 28: 14165–14175, 2008.

Gallivan JP, Logan L, Wolpert DM, Flanagan JR. Parallel specification of competing sensorimotor control policies for alternative action options. Nat Neurosci 19: 320–326, 2016.

Gallivan JP, Stewart BM, Baugh LA, Wolpert DM, Flanagan JR. Rapid automatic motor encoding of competing reach options. Cell Rep 18: 1619–1626, 2017.

Georgopoulos AP, Kalaska JF, Massey JT. Spatial trajectories and reaction times of aimed movements: Effects of practice, uncertainty, and change in target location. J Neurophysiol 46: 725–743, 1981.

Haar S, Donchin O. A revised computational neuroanatomy for motor control. J Cogn Neurosci 32: 1823– 1836, 2020.

Haith AM, Huberdeau DM, Krakauer JW. Hedging your bets: Intermediate movements as optimal behavior in the context of an incomplete decision. PLoS Comput Biol 11: e1004171, 2015.

Haith AM, Pakpoor J, Krakauer JW. Independence of movement preparation and movement initiation. J Neurosci 36: 3007–3015, 2016.

Izawa J, Shadmehr R. On-Line Processing of Uncertain Information in Visuomotor Control. J Neurosci 28: 11360–11368, 2008.

Keyser J, Medendorp WP, Selen LPJ. Task-dependent vestibular feedback responses in reaching. J Neurophysiol 118: 84–92, 2017.

Knill DC, Bondada A, Chhabra M. Flexible, Task-Dependent Use of Sensory Feedback to Control Hand Movements. J Neurosci 31: 1219–1237, 2011.

Lakens D. Calculating and reporting effect sizes to facilitate cumulative science: A practical primer for t-tests and ANOVAs. Front Psychol 4: 1–12, 2013.

Lee JH. Model predictive control: Review of the three decades of development. Int J Control Autom Syst 9: 415–424, 2011.

Li Z, Mazzoni P, Song S, Qian N. A single, continuously applied control policy for modeling reaching movements with and without perturbation. Neural Comput 30: 397–427, 2018.

Lowrey CR, Nashed JY, Scott SH. Rapid and flexible whole body postural responses are evoked from perturbations to the upper limb during goal-directed reaching. J Neurophysiol 117: 1070–1083, 2017.

Mathew J, Crevecoeur F. Adaptive feedback control in human reaching adaptation to force fields. Front Hum Neurosci 15: 742608, 2021.

Mathew J, Lefevre P, Crevecoeur F. Rapid changes in movement representations during human reaching could be preserved in memory for at least 850ms. eneuro 7, 2020.

Mazzoni P, Hristova A, Krakauer JW. Why don’t we move faster ? Parkinson’s disease, movement vigor and implicit motivation. J Neurosci 27: 7105–7116, 2007.

Nashed JY, Crevecoeur F, Scott SH. Influence of the behavioral goal and environmental obstacles on rapid feedback responses. J Neurophysiol 108: 999–1009, 2012.

Nashed JY, Crevecoeur F, Scott SH. Rapid Online Selection between Multiple Motor Plans. J Neurosci 34: 1769–1780, 2014.

Omrani M, Diedrichsen J, Scott SH. Rapid feedback corrections during a bimanual postural task. J Neurophysiol 109: 147–161, 2013.

Oostwoud Wijdenes L, van Beers RobJ, Medendorp WP. Vestibular modulation of visuomotor feedback gains in reaching. J Neurophysiol 122: 947–957, 2019.

Orban de Xivry J-J, Legrain V, Lefèvre P. Overlap of movement planning and movement execution reduces reaction time. J Neurophysiol 117: 117–122, 2017.

Poscente SV, Peters RM, Cashaback JGB, Cluff T. Rapid feedback responses parallel the urgency of voluntary reaching movements. Neuroscience 475: 163–184, 2021.

Prablanc C, Martin O. Automatic control during hand reaching at undetected two-dimensional target displacements. J Neurophysiol 67: 455–469, 1992.

Pruszynski JA, Kurtzer I, Scott SH. Rapid Motor Responses Are Appropriately Tuned to the Metrics of a Visuospatial Task. J Neurophysiol 100: 224–238, 2008.

Pruszynski JA, Scott SH. Optimal feedback control and the long-latency stretch reflex. Exp Brain Res 218: 341–359, 2012.

Sarlegna FR, Mutha PK. The influence of visual target information on the online control of movements. Vision Res 110: 144–154, 2015.

Scholz JP, Schoner G, Latash ML. Identifying the control structure of multijoint coordination during pistol shooting. Exp Brain Res 135: 382–404, 2000.

Scott SH. Optimal feedback control and the neural basis of volitional motor control. Nat Rev Neurosci 5: 532–546, 2004.

Scott SH. A Functional Taxonomy of Bottom-Up Sensory Feedback Processing for Motor Actions. Trends Neurosci 39: 512–526, 2016.

Shadmehr R, Krakauer JW. A computational neuroanatomy for motor control. Exp Brain Res 185: 359– 381, 2008.

Soechting JF, Lacquaniti F. Modification of Trajectory of a Pointing Movement in Response to a Change in Target Location. J Neurophysiol 49: 548–564, 1983.

Todorov E. Optimality principles in sensorimotor control. Nat Neurosci 7: 907–915, 2004.

Todorov E, Jordan MI. Optimal feedback control as a theory of motor coordination. Nat Neurosci 5: 1226–1235, 2002.

Togo S, Yoshioka T, Imamizu H. Control strategy of hand movement depends on target redundancy. Sci Rep 7: 1–7, 2017.

Vetter P, Flash T, Wolpert DM. Planning movement in a simple redundant task. Curr Biol 12: 488–491, 2002.

Wong AL, Haith AM. Motor planning flexibly optimizes performance under uncertainty about task goals. Nat Commun 8: 1–10, 2017.

Wong AL, Haith AM, Krakauer JW. Motor planning. Neuroscientist 21: 385–398, 2015.

